# Prefronto-subcortical hypoconnectivity in schizophrenia: translation of critical pathways for symptom-related functions in nonhuman primates

**DOI:** 10.1101/2023.02.17.528919

**Authors:** Noriaki Yahata, Toshiyuki Hirabayashi, Takafumi Minamimoto

## Abstract

Recent advances in genetic neuromodulation technology have enabled circuit-specific interventions in nonhuman primates (NHPs), thereby revealing the causal functions of specific neural circuits. Going forward, an important step is to use these findings to better understand neuropsychiatric and neurological disorders in humans, in which alterations in functional connectivity between brain regions are demonstrated. We recently identified the causal roles of the pathways from the dorsolateral prefrontal cortex (DLPFC) to the lateral part of the mediodorsal thalamic nucleus (MDl) and dorsal caudate nucleus (dCD) in working memory and decision-making, respectively. In the present study, we examined the resting-state functional connectivity of these two prefronto-subcortical circuits in healthy controls (HCs) and patients with various neuropsychiatric disorders including schizophrenia (SCZ), major depressive disorder (MDD), and autism spectrum disorders (ASD) in humans. We found that the functional connectivity of two pathways, DLPFC-MDl and DLPFC-dCD, was significantly reduced in the SCZ groups compared to HCs; however, this hypoconnectivity was not observed in the ASD or MDD groups, suggesting a disease-specific profile of altered prefronto-subcortical connectivity at rest. These results suggest that causal findings of pathway-specific functions revealed in NHPs can be effectively translated to identify the altered connectivity in neuropsychiatric disorders with related symptoms in humans.

## 1. Introduction

Over the past few decades, comparative studies of brain structure and function between humans and nonhuman primates (NHPs) have been conducted (Petrides and Pandya, 1999; Thiebaut de Schotten et al., 2019). Functional magnetic resonance imaging (fMRI) studies have compared task- and sensory-evoked brain activity (Nakahara et al., 2002; Orban et al., 2004; Wilson et al., 2015) and resting-state functional connectivity (FC) (Hutchison et al., 2012; Sallet et al., 2013) between humans and NHPs, and revealed their shared and distinct functions of specific brain regions or connections, therefore providing a rich insight into human brain evolution. However, because previous studies were conducted on intact brains, the obtained functional implications remained correlative. Beyond correlation, the causal understanding of the higher brain functions of NHPs has been facilitated by recent advancements in neuromodulation techniques. In particular, genetic neuromodulation, such as chemogenetics (Roth, 2016) disturbs specific pairs of anatomically connected brain regions, allowing certain behavior to be causally linked to specific across-region communication (Oyama et al., 2021).

Altered connectivity patterns in the human brain have been demonstrated to be a hallmark of psychiatric and neurodevelopmental disorders. Notably, recent neuroimaging-based data-driven investigations have revealed altered neural networks across the whole brain in many psychiatric disorders. Machine learning (ML) techniques have identified alterations in patterns of resting-state FC that have facilitated diagnostic status prediction in both psychiatric patients and healthy controls (HCs) in schizophrenia (SCZ; Lei et al., 2022; Yoshihara et al., 2020), autism spectrum disorder (ASD; Guo et al., 2017; Kam et al., 2017; Yahata et al., 2016; Yamagata et al., 2019), and major depressive disorder (MDD; Drysdale et al., 2017; Ichikawa et al., 2020; Yamashita et al., 2020). These predictions were shown to be accurate and statistically meaningful, but insufficient for use in clinical settings (e.g., Plitt et al., 2015). This requires the development of methods that enable a more efficient and reliable feature extraction from a large set of neuroimaging data. In addition, the development of methods that facilitate the interpretation of the extracted features is equally important. Without well-documented evidence from previous functional imaging studies, features selected by data-driven procedures cannot always offer a clear connection between the pathophysiology and symptomatology of a disorder. This presents a potential barrier to establishing novel diagnostic and therapeutic tools based on the features discovered; however, owing to the non-invasive nature of human neuroimaging, this cannot be fully resolved solely through human investigation.

Accordingly, the next important step in human-NHP comparative research is to identify the possibility of directly assessing the relationships between a given symptom subserved by altered connectivity in human patients and analogous behavioral changes in NHPs whose connectivity was manipulated. This can be accomplished in two opposite directions (Fig. 1). First, by establishing an NHP model of a given symptom in human disorders, we can understand the causal relationship between FC dysregulation and the symptomology (Fig. 1, from human to NHP). Second, the causal relationship between a given behavioral deficit and anatomically fine-grained connectivity identified in NHPs allows us to elaborate on the anatomical knowledge of the corresponding symptoms in humans (Fig. 1, from NHP to human). In particular, findings from NHP studies can complement data-driven investigations by providing important clues beyond those obtained from human research alone. Owing to the wealth of anatomical and functional knowledge accumulated in subcortical structures and cortico-subcortical connectivity (Aggleton and O’Mara, 2022; Baxter, 2013; Mitchell, 2015), NHP studies may offer an optimal spatial scale at which FC should be evaluated in human studies, thereby enhancing the detectability of disorder-specific alterations.

**Figure 1.**
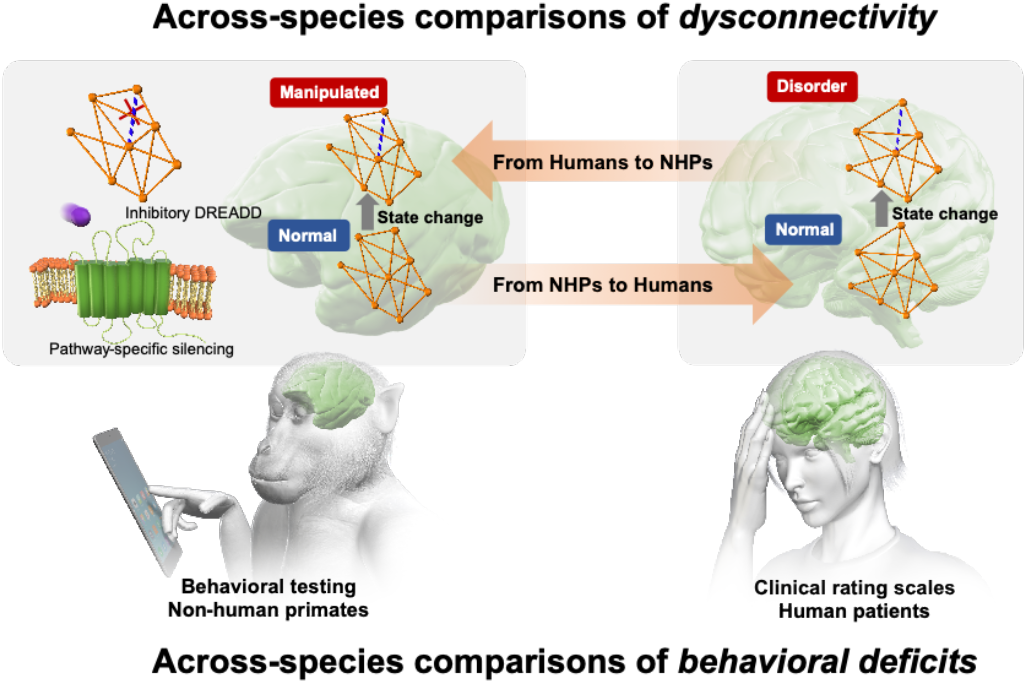
Concept of across species comparisons. Schematic illustration representing two possible comparisons of between humans and NHPs in network dysconnectivity and behavioral deficits. *Top arrow*: An approach from humans to NHPs: Establishing an NHP model by chemogenetically inducing network dysconnectivity related to a given symptom in human disorders. *Bottom arrow:* An approach from NHPs to humans: causal relationships between a given behavioral deficit and anatomically fine-grained connectivity revealed in the NHP study would elaborate on the anatomical knowledge regarding the cause of the corresponding symptoms in humans.

As a key trigger for adopting the second approach, our recent monkey study identified the causal roles of two prefronto-subcortical pathways. Specifically, using imaging-based chemogenetic methods, we selectively silenced the pathways from the dorsolateral prefrontal cortex (DLPFC) to the dorsal caudate (dCD) and lateral mediodorsal thalamus (MDl) individually. We demonstrated these pathways’ dissociable roles in cognitive functions, which are critical for working memory (WM) and decision-making (Oyama et al., 2021) (Fig. 2A). In most human neuroimaging studies, however, the thalamus has not been investigated at this fine level of regional parcellation; it is usually defined as a single region or set of coarsely segregated regions that depended on image-based delineation. Although aberrant patterns of thalamocortical connectivity have been reported in various psychological disorders including SCZ, MDD, and ASD (Woodward, 2017), the exact involvement of the encompassing nuclei remains largely unidentified. This necessitates the extension of the causal relationships obtained with NHPs to the pathophysiology of human neuropsychiatric disorders, with special considerations to thalamic nuclei.

**Figure 2.**
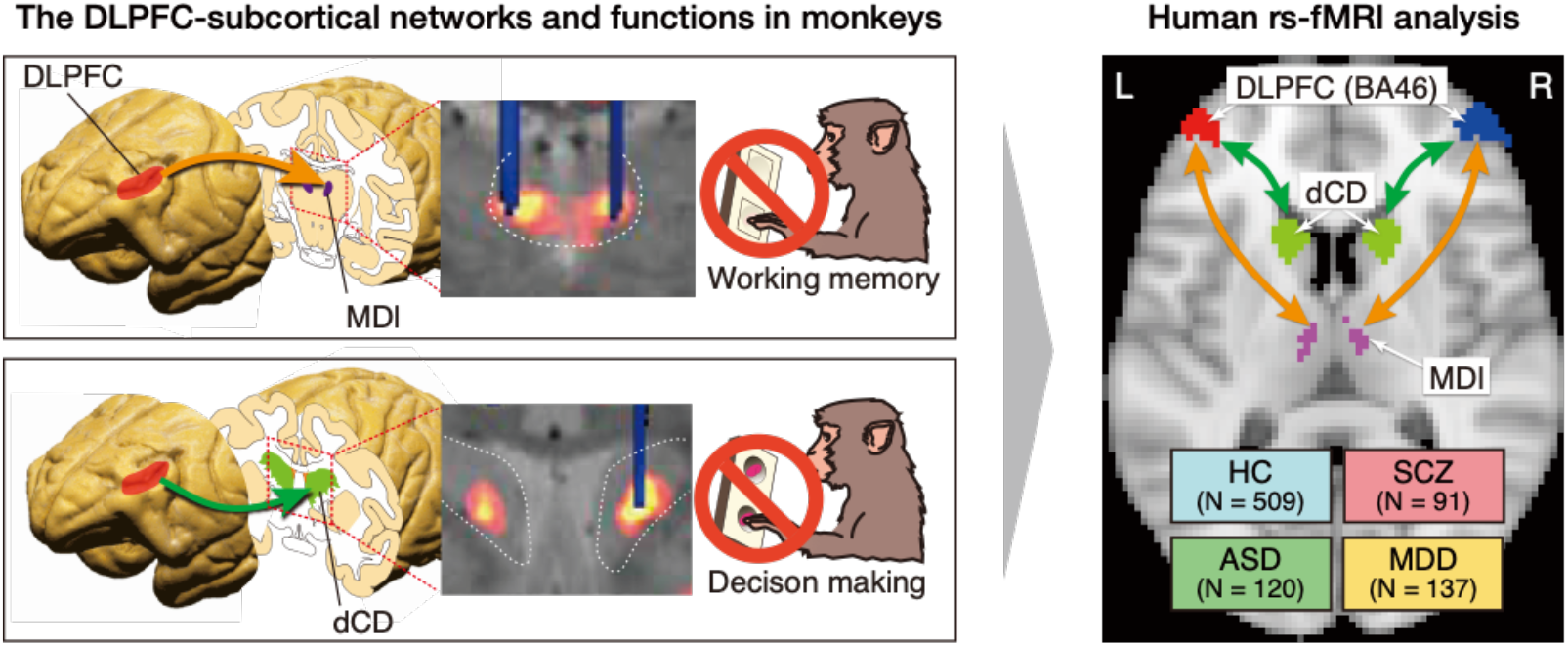
Design of this study. **A**. A schematic illustration representing the chemogenetic pathway-selective silencing of prefronto-subcortical pathway. Introduction of an inhibitory designer receptor hM4Di into neurons in the DLPFC led to expression of hM4Di in axon terminals in the projection sites, the MDl and dCD. The middle panel shows hM4Di expression at the projection site visualized by PET (hot color) and DREADD agonist injection through the cannula visualized by CT (blue). Synaptic silencing induced by agonist infusion impaired WM and decision-making, respectively. Modified from (Oyama et al., 2021). **B**. An illustration representing the voxels of interest for DLPFC, dCD, and MDl in human fMRI data. Abbreviations: DLPFC, dorsolateral prefrontal cortex; MDl, mediodorsal thalamic nucleus; dCD, dorsal caudate nucleus; WM, working memory; HC, healthy control; SCZ, schizophrenia; MDD, major depressive disorder; ASD, autism spectrum disorder.

In the present study, we examined the resting-state FC of these two prefronto-subcortical pathways in HCs and three groups of patients with SCZ, MDD, and ASD (Fig. 2B). We found that FC of the two pathways, DLPFC-MDl and DLPFC-dCD, was significantly and specifically reduced in the SCZ group compared to HCs. Our results suggest that the pathway-specific function causally revealed in NHPs can be used to elucidate the relationship between altered FC and symptoms in human psychiatric disorders.

## 2. Materials and Methods

### 2.1. Datasets

The current study was based on two datasets from a multi-site, multi-disorder neuroimaging database compiled by the Strategic Research Program for Brain Sciences (SRPBS) supported by the Japan Agency for Medical Research and Development (AMED).

First, we used the SRPBS Multi-disorder MRI Dataset (Dataset 1; available at https://bicr-resource.atr.in/srpbsopen/) to evaluate interregional FC among patient groups and HCs. Dataset 1 was a collection of anonymized 3T MR images and associated demographic/clinical data of over 1,400 participants recruited at 11 imaging sites in Japan. Each participant underwent an MR scan at one imaging site to acquire resting-state functional and T_1_-weighted structural MR images. A B_0_ field map was also available for the majority of participants. In the present analysis, we incorporated data with acquisition conditions compatible with the unified imaging protocol determined in the SRPBS MRI guidelines (Tanaka et al., 2021). This led us to form a local patient dataset that included participants and HCs from six sites (site codes = ATT, ATV, COI, KUT, SWA, and UTO; Table 1). Table 1 summarizes the demographic properties of the local patient dataset.

**Table 1.** Summary of the participants in the local patient dataset.

| Group | Age<br>(years) | $N$ | Sex<br>(male/female) | Site | | | | | |
| --- | --- | --- | --- | --- | --- | --- | --- | --- | --- |
|  |  |  |  | ATT | ATV | COI | KUT | SWA | UTO |
| HC | $38.9 \pm 15.7$ | 509 | (288/221) | X | X | X | X | X | X |
| SCZ | $37.9 \pm 11.3$ | 91 | (56/35) | | | X | X | X | |
| ASD | $32.4 \pm 7.8$ | 120 | (104/16) | | | | | X | X |
| MDD | $41.7 \pm 12.5$ | 137 | (73/64) | | X | X | | X | |
| Total | $38.3 \pm 14.2$ | 857 | (521/336) | | | | | | |

Abbreviations: ASD = autism spectrum disorder, HC = healthy controls, MDD = major depressive disorder, SCZ = schizophrenia.

Second, we used the SRPBS Traveling Subject MRI Dataset (Dataset 2; available at https://bicr-resource.atr.jp/srpbsts/), for which nine healthy participants visited multiple imaging sites including the six sites in Dataset 1 (hence the name “traveling subjects”). They underwent functional and structural MR scans that conformed to the unified imaging protocol in the SRPBS MRI guidelines (Tanaka et al., 2021). In general, multisite data are subject to both measurement and sampling biases. Measurement bias refers to differences in the instrumental and imaging properties of MR data acquisition, whereas sampling bias refers to site-to-site differences in the composition of participant groups (Yamashita et al., 2019). The presence of these two biases complicates across-site comparison and interpretation of the results. Data acquired from traveling subjects can help mitigate this problem because they quantitatively describe how measurement bias impacts imaging results within the same individual. Previously, we combined Datasets 1 and 2 to develop an analytical method that enabled the optimal reduction of measurement bias, thereby achieving “harmonization” of the multisite resting-state fMRI data (Yamashita et al., 2019). Here, we employed an equivalent harmonization scheme to the local patient dataset to enable a comparison of FC across sites (see section 2.4). For this purpose, we formed a local traveling subject dataset consisting of data from eight sites in Dataset 2 (site codes = ATT, ATV, COI, HKH, HUH, KUT, SWA, and UTO). At these sites, the manufacturers of the MR systems were either GE (Chicago, IL, USA) or Siemens (Munich, Germany), which coincided with the local patient dataset.

### 2.2. Data preprocessing

To preprocess and denoise the raw data and calculate the mean time course within the regions of interest, we used the CONN Toolbox (version 19c; https://web.conn-toolbox.org/) running SPM12 in MATLAB (version R2018a; MathWorks, Natick, MA, USA). We used a built-in default preprocessing pipeline that corrects for head motion (realignment), unwarping, slice-timing correction, and spatial normalization of the images to the Montreal Neurological Institute space. To reduce the physiological and non-neuronal noise present in the temporal fluctuation of blood oxygenation level-dependent (BOLD) signals, we then performed subject-level denoising. Specifically, we used a regression model that incorporated the following confounders: (1) six motion parameters, their first-order derivatives, and their quadratic effects (24 parameters in total); (2) five principal components calculated in each of the white matter and cerebrospinal fluid; and (3) excessive frame-to-frame head motions in the form of a binary vector indicating the numbers of scans. CONN’s built-in algorithm (ArtRepair) was used to evaluate head motion with conservative thresholds (global signal: z-value = 3; subject motion: 0.5 mm). The residual was then band-pass filtered (transmission range: 0.008–0.09 Hz), yielding a noise-reduced BOLD time course for the subsequent calculation of FCs.

### 2.3. Definition of regions of interest and calculation of interregional FC

The spatial extent of anatomical regions in the brain was determined by using the Automated Anatomical Labelling atlas 3 (AAL3) (Tian et al., 2020). Regarding the three regions of interest in the present study (DLPFC, MDl, and dCD), we made the following modifications so that their boundaries better corresponded to those investigated in our previous macaque studies (Fig. 2). The DLPFC was defined as the Brodmann area (BA) 46 of the middle frontal gyrus (MFG). We combined the AAL3 and Brodmann maps (available in MRIcron software; (Rorden and Brett, 2000)) to demarcate the BA 46 boundary in the MFG. AAL3 was also used to define MDl; however, because the parcellation of the caudate nucleus in the AAL3 does not allow isolation of the dCD, we used an alternative parcellation of the human subcortex reported elsewhere (Tian et al., 2020). As in our previous macaque study (Oyama et al., 2021), we did not consider the functional lateralization of the subcortex structures so that both the MDl and dCD encompassed their counterparts in the left and right hemispheres. Conversely, given the functional lateralization in the human prefrontal cortex, parts of the DLPFC in the left and right hemispheres were treated separately.

The boundaries of other regions were adopted from the AAL3, except for the cerebellum, wherein their subregions were reorganized into three regions: the left and right cerebellum and vermis. This was to consider the inconsistent imaging coverages of the cerebellum among participants, which could lead to missing elements in their interregional correlation matrices.

Overall, 119 brain regions were identified in the present study. The mean time course was calculated for each region using normalized, denoised functional images. Finally, an interregional correlation matrix was formed for each participant by exhaustively calculating the Pearson correlations for all pairs of regions. The FCs between the DLPFC (left[L]/right[R]) and MDl and between the DLPFC(L/R) and dCD were the targets of the present investigation, whereas the harmonization procedure described in the next section required FCs across the brain.

### 2.4. Reduction of measurement bias and confounding effects by age and sex

The correlation matrices of all subjects were submitted to a procedure to reduce: (1) measurement bias, and (2) the effects of age and sex in the present FC data. The methodological details to reduce measurement bias have been reported previously (Yamashita et al., 2019). Briefly, for FC in the *k*^th^ participant (where *k* runs over all participants), we constructed a regression model of the form:

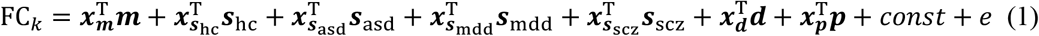

Column vectors ***m***, ***s***_hc_,_asd_,_mdd_,_scz_, ***d*** and ***p*** represent the measurement bias (six sites × 1), sampling bias of HC (six sites × 1), ASD (two sites × 1), MDD (three sites × 1), SCZ (three sites × 1), disorder factor (three disorder types × 1), and identity of a traveling subject if applicable (nine subjects × 1). T denotes the transposition of the column vector. The ***x***-vectors in (1) describe the following properties of the *k*^th^ participant: the imaging site (***x_m_***), disorder type (***x_d_***), site information per disorder (***x***_***s***hc,asd,mdd,scz_), and the identity of the traveling subjects (***x_p_***). In each vector, the element applicable to the *k*^th^ participant is set to 1, and to 0 otherwise. Finally, *e* denotes the residual of the model. Given the set of equations (1) derived from all participants, we solved for ***m***, _***s***hc,asd,mdd,scz_, ***d***, ***p***. (see (Yamashita et al., 2019) for details). The measurement bias-corrected FC’ is calculated as follows:

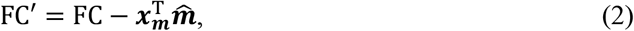

where 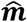 is the estimated measurement bias.

Furthermore, to consider group differences in age and sex compositions in the present dataset (Table 1), we extended regression model (1) to incorporate their effects:

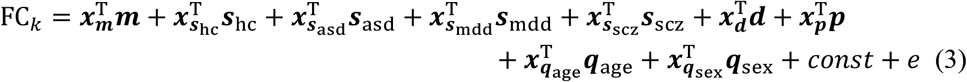

where ***q***_age,sex_ and ***x***_***q***age,sex_ are the linear effects and the participants’ value of normalized age and sex (binary), respectively. Solving the set of equations (3) of all the participants, and using the estimates 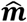, 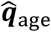, and 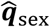, we obtained FC values corrected for measurement bias and the effects of age and sex:

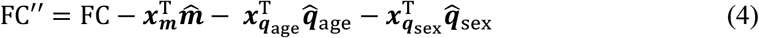

We refer to this FC” as FC in the subsequent analyses.

### 2.5. Evaluation of FC among regions of interest

Here, we focused on four FC estimates connecting (1) the DLPFC(L) and MDl, (2) DLPFC(R) and MDl, (3) DLPFC(L) and dCD, and (4) DLPFC(R) and dCD (Fig. 2B). For each FC estimate, we conducted a one-way analysis of variance (ANOVA) to evaluate group differences in the means (HC, SCZ, ASD, and MDD) at a statistical threshold of *P* = 0.05. If a significant effect of group was detected, we further performed post-hoc *t*-tests to identify the pair(s) of groups exhibiting a significant difference in FCs. The Tukey-Kramer procedure was used to correct for multiple comparisons.

In SCZ participants, for the FC values that exhibited statistically significant deviations from HCs, we examined whether their FC values were correlated with their clinical profiles. Specifically, we evaluated the memory performance of the patients using verbal memory and WM measures from the Brief Assessment of Cognition in Schizophrenia (Kaneda et al., 2007). We also evaluated the effect of medication on FC in SCZ using the chlorpromazine-equivalent dose (mg) of antipsychotics administered to the patients.

## 3. Results

### 3.1. Hypoconnectivity of the DLPFC-MDl and DLPFC-dCD pathways in SCZ

For all four pathways considered, one-way ANOVA revealed a significant group effect on FC means (*P* < 0.05). Post-hoc *t*-tests revealed that the SCZ group exhibited significant hypoconnectivity compared with the HC group (corrected *P* = 0.023, 0.019, 0.005, and 0.024 for the FC values of DLPFC(L)-MDl, DLPFC(R)-MDl, DLPFC(L)-dCD, and DLPFC(R)-dCD, respectively; see Fig. 3AB) and ASD (corrected *P* = 0.048 for the FC value of DLPFC(L)-dCD). In other words, all detected group differences were related to the hypoconnectivity of the SCZ group.

**Figure 3.**
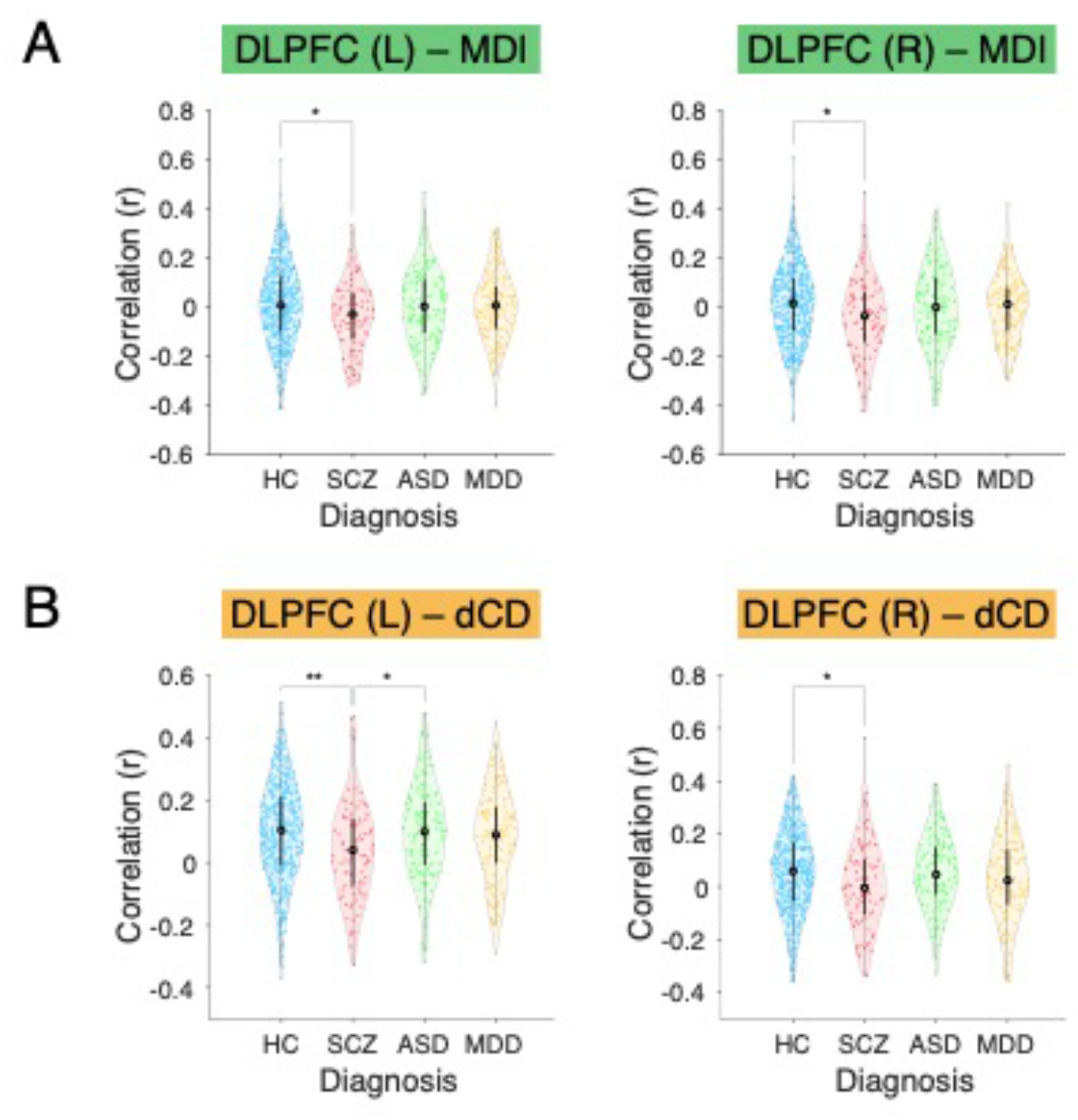
Reduced FC of the DLPFC–subcortical pathways in patients with schizophrenia. Violin plots represent distributions of correlation coefficients of the mean BOLD signal time courses between the DLPLC-MDl (A) and the DLPFC-dCD (B). The left and right DLPFC to each subcortical region were analyzed separately and are shown in left and right, respectively. Dots represent individual measurements, and the shape of the plots represents the data distribution. The asterisk represents a significant difference at * *P* < 0.05 and ** *P* < 0.01. Abbreviations: DLPFC, dorsolateral prefrontal cortex; MDl, mediodorsal thalamic nucleus; dCD, dorsal caudate nucleus; BOLD, blood oxygenation level-dependent; HC, healthy control; SCZ, schizophrenia; ASD, autism spectrum disorder; MDD, major depressive disorder.

### 3.2. No direct correlation between DLPFC-subcortical connectivity and memory scores in SCZ

WM impairment and related dysfunction are among the best-replicated endophenotypes in SCZ. Indeed, the scores of the verbal memory and WM tests were significantly lower in the SCZ group than the HC group (two-sample *t*-tests, *P* = 8.9 × 10^-10^ and 6.0 × 10^-5^, respectively; Fig. 4A and C). This observation questions whether there is a relationship between reduced DLPFC and subcortical structure connectivity and WM impairments in SCZ; however, as shown in Figs. 4B and D, there was no significant relationship between FC and memory scores for either DLPFC-subcortical pathway.

**Figure. 4.**
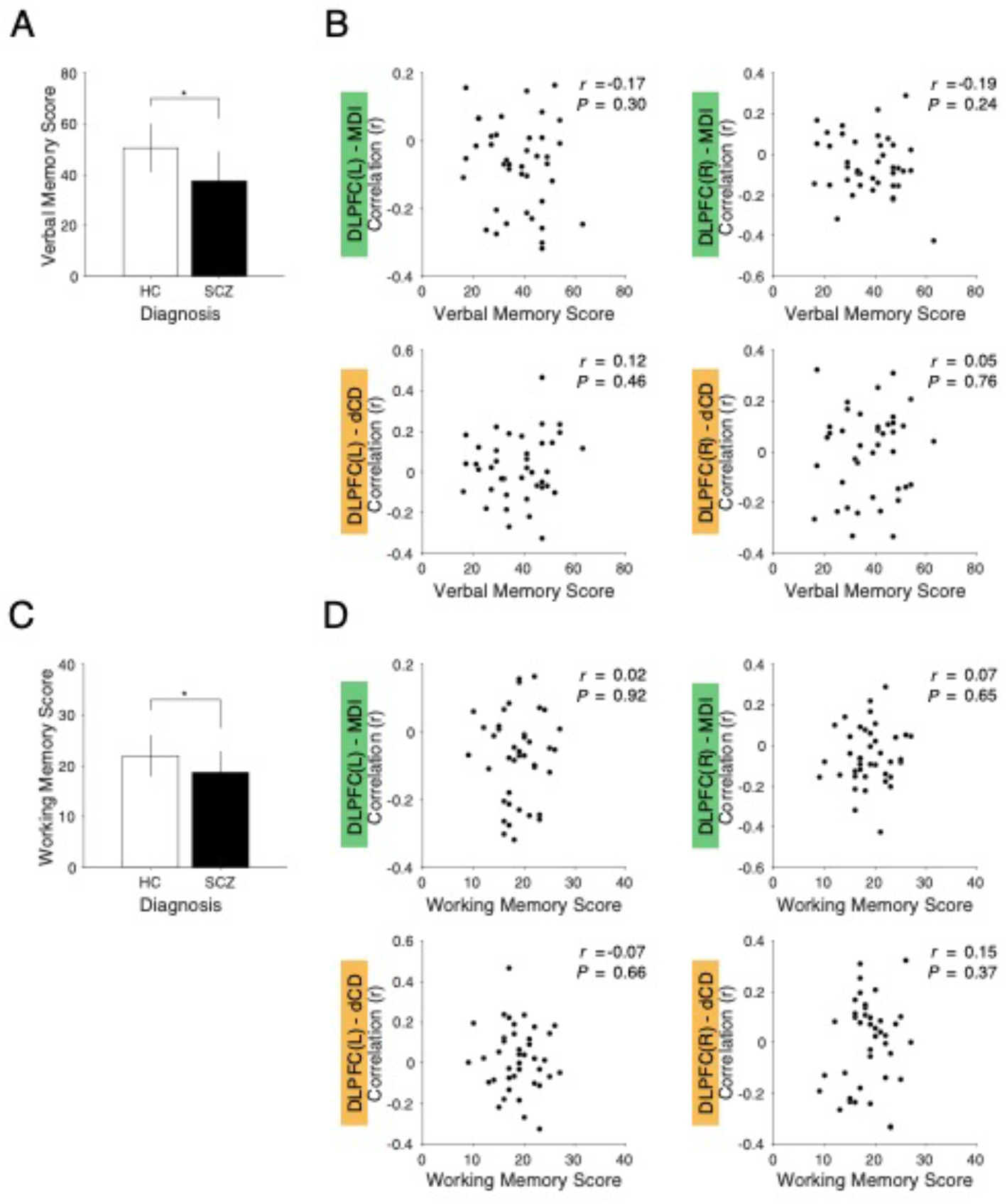
No direct relationships between FC of the DLPFC–subcortical pathways and memory-related test performance in the SCZ group (*N* = 40). **A.** Verbal memory scores (mean ± standard deviations [s.d.]) in the HC and SCZ groups. **B**. Relationships between FC and memory test scores for left DLPLC-MDl (upper left), right DLPLC-MDl (upper right), left DLPFC-dCD (lower left), and right DLPFC-dCD pathways (lower right). **C**. WM scores (mean ± s.d.) in the HC and SCZ groups. **D**. Relationships between FC and memory scores. Asterisks indicate a significant difference (*P* < 10^-5^). Abbreviations: FC, functional connectivity; DLPFC, dorsolateral prefrontal cortex; HC, healthy control; SCZ, schizophrenia; MDl, mediodorsal thalamic nucleus; dCD, dorsal caudate nucleus; WM, working memory.

The observed hypoconnectivity in SCZ may be related to the administration of antipsychotics, such as dopamine D_2_ receptor antagonists. To evaluate this possibility, we used the chlorpromazine-equivalent dose (mg) of antipsychotics administered to the patients in the patient dataset (*N* = 40). A significant negative correlation was found between the magnitude of DLPFC-dCD FC and the dosage of antipsychotic medication (*r* = −0.32, *P* = 0.042; Fig. 5).

**Figure 5.**
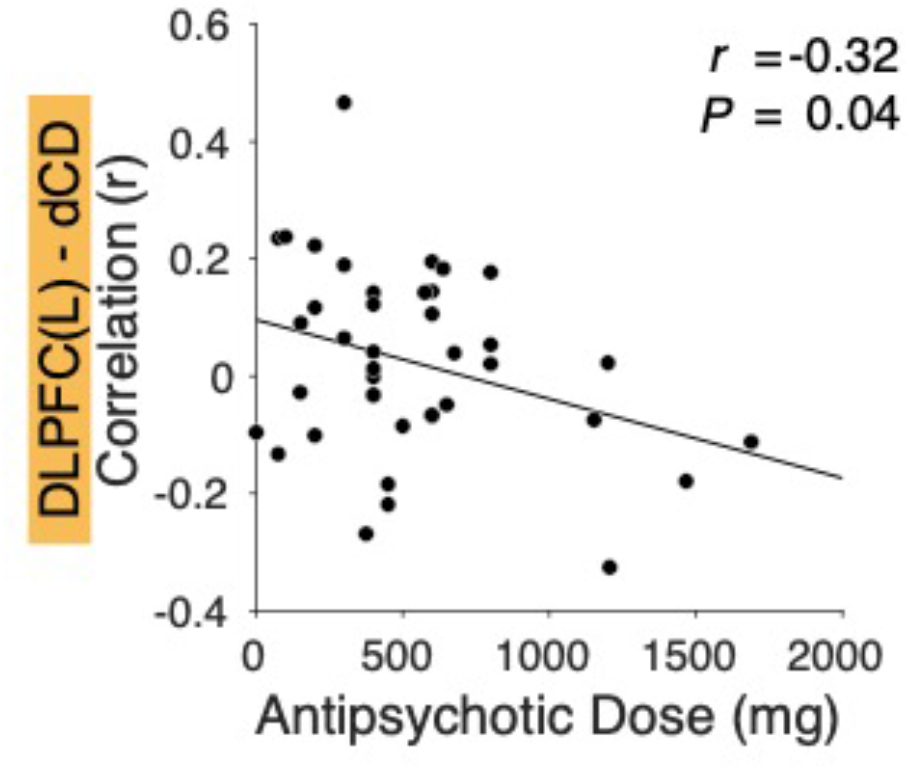
Significant negative correlation between the dose of antipsychotic medication and the FC of left DLPFC-dCD (Pearson correlation, *r* = −0.32, *P* = 0.042). Finally, we confirmed that no significant correlation was found between the strength of the DLPFC-dCD and DLPFC-MDl connections in either hemisphere in the SCZ group (Fig. 6). This suggests that the observed reduced connectivity in the SCZ group does not simply reflect a general dysfunction of the prefronto-basal ganglia-thalamocortical loop.

**Figure 6.**
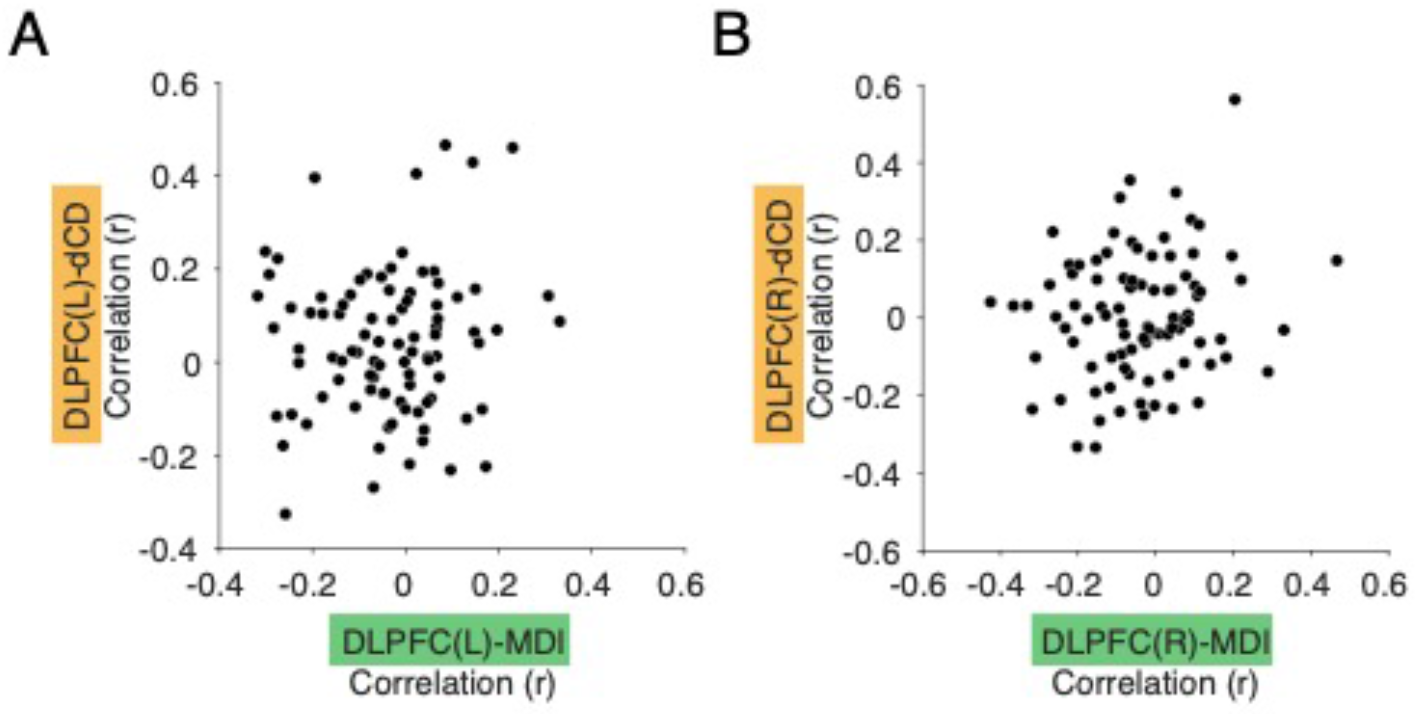
No significant relationship between DLPFC-dCD and DLPFC-MDl FC in the SCZ group.

## 4. Discussion

In the present study, we performed a translation of the causal evidence of NHPs circuit manipulation into humans with disorders characterized by network dysconnectivity and related behavioral deficits. We used resting-state fMRI data from patients with one of three disorders (SCZ, ASD, and MDD) and HCs to assess FC among human homologous regions in the prefronto-subcortical pathways. Compared to HCs, SCZ patients showed reduced FC in the DLPFC-MDl and DLPFC-dCD networks. Critically, this hypoconnectivity was not observed in patients with ASD or MDD, suggesting a disease-specific profile.

Numerous resting-state fMRI studies have consistently reported prefronto-thalamic hypoconnectivity in SCZ patients, which has been implicated as the pathophysiology of this disease (Anticevic et al., 2014; Anticevic et al., 2015; Avram et al., 2018; Cheng et al., 2015; Tu et al., 2015; Woodward and Heckers, 2016; Woodward et al., 2012). Likewise, hypoconnectivity between the PFC and the dCD has also been reported in SCZ patients and individuals at risk for psychosis (Dandash et al., 2014; Fornito et al., 2013; Zhou et al., 2007). In these studies, changes in FC were assessed either exploratively using seed-based analysis with coarsely predetermined regions of interest in the thalamus, striatum, or cortex, or using a data-driven approach such as voxel-based analysis. Therefore, for the prefronto-thalamic hypoconnectivity in SCZ, the exact loci of the thalamus and cortex differed across studies, making it difficult to determine its relevance with the symptoms and pathophysiology of SCZ.

Moreover, comparing changes in FC across multiple diseases would complicate the analysis and require more data, hindering the investigation of disease or symptom specificity. In contrast, we adopted an “NHP-to-human approach” by applying human homolog circuits of the pathways whose functions were causally identified in NHPs (Fig. 2), revealing specific hypoconnectivity to the SCZ group (Fig. 3). These disease-specific profiles of circuit alterations can be discussed alongside the critical functions of the DLPFC-MDl and DLPFC-dCD pathways revealed in NHPs, namely WM and decision-making.

Despite the significant reductions in memory-related abilities in the SCZ group, individual differences in FC in the DLPFC-subcortical network did not account for these memory disturbances (Fig. 4). This was also the case in previous resting-state fMRI studies with SCZ patients; the relationship between individual differences in thalamocortical dysconnectivity and executive cognitive abilities has been inconsistent and controversial (Giraldo-Chica and Woodward, 2017; Woodward, 2017). One interpretation of this discrepancy may be due to the heterogeneity of this disease and/or patient samples, simply reflecting the complex and broader dysfunction that characterizes the disease. Other plausible explanations include mismatches between humans and NHPs, specifically regarding how cognitive abilities and connectivity were measured. First, WM scores in the SCZ group typically target verbal memory, whereas the monkey circuit manipulation referred to in the present study focused on spatial WM. Second, alterations in FC at rest may not necessarily reflect those during the task. Indeed, recent fMRI studies during task emphasized the relationship between PFC-thalamic hypoconnectivity and cognitive disabilities in SCZ (Huang et al., 2019; Wu et al., 2022). Future studies should examine the connectivity in SCZ patients during spatial WM tasks. In addition to WM, dysregulation of decision-making has also been reported in SCZ. However, no behavioral assessment related to decision-making was available in the present dataset, hence we did not address the function of DLPFC-dCD in the SCZ patients. Importantly, striatal-cortical dysconnectivity has been reported to be related to dopamine dysregulation in unmedicated SCZ patients (Horga et al., 2016). As dysconnectivity of DLPFC-dCD correlated with anti-psychotic medication (Fig. 5), future studies should investigate the alterations in the DLPFC-dCD pathway and decision-making, preferably in unmedicated SCZ patients.

As mentioned above, aberrant thalamocortical FC has been repeatedly observed in previous neuroimaging studies of SCZ, but the exact involvement of specific thalamic nuclei remains undetermined owing to the inconsistent anatomical demarcation adopted. We utilized the knowledge from an NHP study and revealed that FC between the MDl and DLPFC was downregulated only in the SCZ group and was intact in both MDD and ASD groups, indicating a successful application of NHP findings in advancing the FC-based investigation of psychiatric disorders. NHP studies can also suggest an optimal “granularity” at which to parcellate the human brain, thereby providing useful information, especially when evaluating interregional FCs, regarding whether a given region may be treated as a functionally homogeneous entity or an aggregate of inhomogeneous subregions. In imaging studies, the choice of the anatomical atlas critically affects the representation of functional networks in the human brain. For example, in our previous ML study that identified 15 FCs as being most relevant in the classification of individuals with SCZ and HCs (Yoshihara et al., 2020), the thalamus was defined as a single anatomical structure, wherein the activities of the individual nuclei were all averaged out. Given that the current study revealed the SCZ-specific reduction of the FC between MDl and DLPFC, future ML investigations should parcellate the thalamus more finely to improve the precision of SCZ-specific feature selection. Accumulation of findings from NHP studies will also provide optimal parcellations in key regions across the whole brain, which is also beneficial in ML-based feature extraction studies. When the derived model overfits to the data, the prediction of a diagnostic label in external data catastrophically fails (Whelan and Garavan, 2014). This can be circumvented by balancing the number of independent variables in the model (i.e., the number of FC values) and sample data (i.e., the number of participants). In most neuroimaging studies, the former typically outnumber the latter, and a reduction of independent variables such as the FCs is required. While the naïve use of coarse parcellation reduces the number of regions to calculate FC estimates, knowledge from NHP studies may prevent the risk of obscuring the disorder/symptom-specific features in FC values. Moreover, the altered FCs selected through data-driven investigations are not necessarily accompanied by interpretations of how they contribute to disorder symptoms. NHP studies with circuit manipulation provide useful information about the relationship between altered FCs and symptoms, facilitating the identification of efficient targets in neurofeedback therapy (Thibault et al., 2018).

In the present study, we found that the DLPFC-MDl connectivity, previously identified as being essential for WM in NHPs, significantly decreased specifically in the SCZ group with decreased WM performance, suggesting the importance of proper parcellation of subcortical brain regions to correctly evaluate disease-related alteration of connectivity (Klein-Flugge et al., 2022). Unlike previous comparative studies on intact brain structures and functions between humans and NHPs (Rocchi et al., 2021; Thiebaut de Schotten et al., 2019), the present results link the causal assessment on the function of a fine-grained connectivity in NHPs with the corresponding pathological dysconnectivity in human patients. Our results provide a new avenue for causal translational research between humans and NHPs, and re-emphasize the importance of macaque research with causal manipulation of a targeted functional circuit. A modest correspondence of the DLPFC between humans and macaques has been demonstrated in terms of both cytoarchitecture (Petrides and Pandya, 1999) and resting-state fMRI connectivity (Sallet et al., 2013). Meanwhile, functional and anatomical knowledge of the MD has been accumulated in macaques, including its subdivisions and their distinct connectivity patterns with the cortex (Baxter, 2013; Mitchell, 2015; Oyama et al., 2021; Phillips et al., 2019), but their relevance and correspondence with the human MD remains largely unestablished, despite recent neuroanatomical study illustrating MD subdivisions and connectivity (Li et al., 2022). The present study thus uniquely provides an invaluable translation of the knowledge of an MD subregion from macaques to humans. Causal translational research in clinical conditions has been successfully conducted with rodents especially for subcortical regions (Low et al., 2021; Zhou et al., 2019). However, the pathophysiology of psychiatric and developmental disorders often includes malfunction of the prefrontal cortex, for which modeling in rodents is challenging (Laubach et al., 2018; Roberts and Clarke, 2019). Therefore, the present approach of causal translational research with NHPs for psychiatric and developmental disorders would provide invaluable opportunities for a deeper understanding of the pathophysiology and for identifying more effective therapeutic targets for such disorders.

Although genetic methods allow the manipulation of specific connectivity in macaques such as from the DLPFC to the MDl, even such focal manipulation would result in more large-scale changes in the activity and/or connectivity (Gerits et al., 2012; Grayson et al., 2016; Hirabayashi et al., 2021; Vancraeyenest et al., 2020). Whole-brain resting-state fMRI investigations in NHPs would thus be an important next step for future research to delineate the complete picture of circuit-manipulation effect in the macaque brain before comparing with disease-related dysconnectivity in human patients. In addition, developing a database of macaque research that summarizes the causal evidence of functional localization would facilitate translational research via the currently proposed approach in the future. Conversely, by creating an NHP model based on a set of connectivity changes related to a specific symptom obtained from human clinical data, it would allow testing the causal role of such connectivity in the disease, leading to a better understanding of the pathophysiology.

In summary, recent advances in genetic neuromodulation technologies provide unique opportunities for comparisons between humans and NHPs in terms of network dysconnectivity and behavioral deficits. Using the NHP-to-human approach, the current study demonstrated that the link between function and fine-grained connectivity in NHPs is useful for investigating the corresponding pathological dysconnectivity in humans. Future translational evidence through circuit manipulation in NHPs will provide a valuable opportunity to better understand the pathophysiology and to develop more effective therapeutic targets.

## Abbreviations

ASD: autism spectrum disorder
dCD: dorsal caudate nucleus
DLPFC: dorsolateral prefrontal cortex
FC: functional connectivity
HC: healthy control
MDD: major depressive disorder
MDl: mediodorsal thalamic nucleus
ML: machine learning
NHP: nonhuman primate
SCZ: schizophrenia

## Data accessibility statement

Data are available upon substantiated request to the corresponding authors.

## CRediT author statement

**Noriaki Yahata:** Conceptualization, Formal analysis, Writing – original draft, Review & Editing, Visualization. **Toshiyuki Hirabayashi:** Conceptualization, Writing – original draft, Review & Editing, Visualization, Funding acquisition. **Takafumi Minamimoto:** Conceptualization, Writing – original draft, Review & Editing, Visualization, Project administration.

## Acknowledgments

We thank K. Yamashita and A. Yamashita for their technical assistance. This study was supported by MEXT/JSPS KAKENHI Grant Numbers JP20H05955 (to TM), and by AMED Grant Numbers JP22dm0307008 (to NY) and JP22dm0307007 (to TH).

